# Rat movements reflect internal decision dynamics in an evidence accumulation task

**DOI:** 10.1101/2023.09.11.556575

**Authors:** Gary A. Kane, Ryan A. Senne, Benjamin B. Scott

## Abstract

Perceptual decision-making involves multiple cognitive processes, including accumulation of sensory evidence, planning, and executing a motor action. How these processes are intertwined is unclear; some models assume that decision-related processes precede motor execution, whereas others propose that movements reflecting on-going decision processes occur before commitment to a choice. Here we develop and apply two complementary methods to study the relationship between decision processes and the movements leading up to a choice. The first is a free response pulse-based evidence accumulation task, in which stimuli continue until choice is reported. The second is a motion-based drift diffusion model (mDDM), in which movement variables from video pose estimation constrain decision parameters on a trial-by-trial basis. We find the mDDM provides a better model fit to rats’ decisions in the free response accumulation task than traditional DDM models. Interestingly, on each trial we observed a period of time, prior to choice, that was characterized by head immobility. The length of this period was positively correlated with the rats’ decision bounds and stimuli presented during this period had the greatest impact on choice. Together these results support a model in which internal decision dynamics are reflected in movements and demonstrate that inclusion of movement parameters improves the performance of diffusion-to-bound decision models.

**Highlights:**

- Development and validation of a free response pulse-based accumulation task for rats
- Response times are well described by drift diffusion models
- Incorporating movement data into diffusion models improves inference of latent decisions variables
- Rats weight sensory evidence most strongly prior to movement

## INTRODUCTION

Perceptual decision-making is a complex and integral part of our interaction with the world. It involves the accumulation of sensory evidence, comparison with internal models, and subsequent decision commitment (Gold and Shadlen 2007). This process allows us to select the appropriate motor response based on the evidence available. Traditional models conceptualize perceptual decision making as a serial process in which evidence accumulation precedes decision commitment and motor execution (Hanks and Somerfield 2017). More recent studies suggest an alternative model in which evidence accumulation and motor actions occur in parallel (Pinto et al., 2018; Musall et al., 2019). In this model, aspects of ongoing motor activity may reflect underlying cognitive dynamics (Shadlen and Kiani, 2013; Wipinski, 2020). Reconciling these models has been challenging and the relationship between motor actions and cognitive processing remains unclear (Lepora and Pezzulo 2015; Selen et al. 2012).

To better understand how movement is related to the decision making process, we developed and applied two complementary methods. The first is a novel free response evidence accumulation task for rodents. In this task, inspired by previous pulsed based accumulation tasks (Brunton et al. 2013; Scott et al. 2015; Gupta et al. 2023) agents observe a series of flashes from two choice ports. Selection of the choice port associated with the higher flash probability was rewarded with a drop of sugar water. This task has two key new features: rats are free to move at any time of the cue period, i.e. no head or nose fixation is involved, and evidence is presented from trial initiation until a choice is reported. This design allowed us to characterize animal movements and evidence accumulation throughout the duration of a trial and to study how movements and decision processes co-occur across the entire trial. The second method movement-based drift diffusion model (DDM) which incorporates pose estimation to constrain decision parameters on a trial-by-trial basis. In this method, termed mDDM, neural network-based video pose estimation is used to track the movements of a decision-maker moment-by-moment. These movement parameters are then used to constrain decision parameters of the DDM on a trial-by-trial basis.

We trained male and female rats to perform the free response task in an automated operant training facility. Rats learned the task quickly, exhibited high accuracy and showed hallmarks of evidence accumulation, including response time distributions that are well fit by a traditional DDM. In parallel, we recorded behavioral videos of rats performing the task and then fit the mDDM. Using this approach, we find that the mDDM outperforms traditional DDMs, based on standard model comparison metrics, and provides a trial-by-trial estimate of decision parameters including starting point, non-decision time and decision threshold. Interestingly we also observe a period of relative immobility, prior to movement onset, and that evidence presented during this period was a better predictor of subsequent choice than later evidence. Together these results are consistent with a model of decision making in which movements reflect the internal dynamics of the decision process, but that there exists a period of deliberation, prior to movements, when the majority of evidence is accumulated. In addition, the task and modeling approach we describe is well-suited for studying the decision-making process and could be paired with neural recordings in future studies to further characterize perceptual decision making in rats and other species.

## RESULTS

### Rats accumulate evidence in a free-response perceptual decision-making task

We developed a free-response version of a visual pulse-based evidence accumulation task (Scott et al. 2015; 2017; 2018) which rats can perform in a 3-port operant chamber (**Figure 1A**). To initiate the trial, the rat would nose poke in the center port. Following trial initiation, lights on the left and right side reward ports flashed bilaterally indicating the start of the cue period.

**Figure 1.**
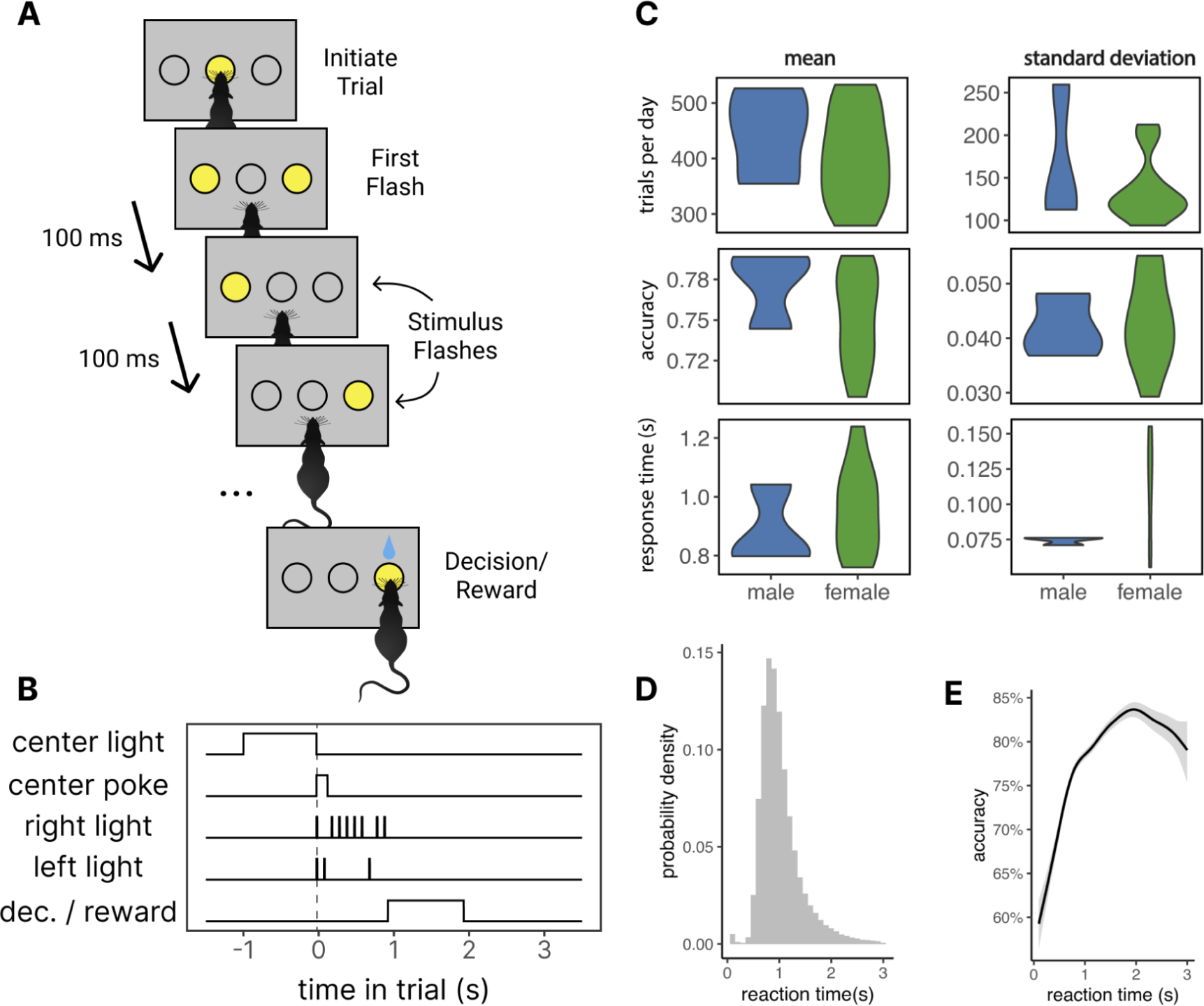
A free response pulse based accumulation task for rats. A) Schematic of the behavioral task. Rats initiate a trial by inserting their nose into the center port of a 3-port operant training chamber. After initiation rats are presented with a series of flashes from the left and right ports according to a Bernoulli process. The trial ends when rats insert their noise into one of the two side ports. Rats obtain reward if they choose the side with the higher flash probability. B) Timing of events in an example behavioral trial. C) Behavioral performance of rats (n = 3 male, 10 female) performing the task. Rats exhibit similar trial numbers (One-way RM ANOVA, p=0.458, F=0.592) (upper panels), accuracy (One-way RM ANOVA, p=0.469, F=0.564) and response time (One-way RM ANOVA, p=0.393, F=0.789) (lower panels) across sexes D) Histogram of rat response times across all rats. E) Accuracy vs. response time curve derived from a generalized additive mixed model.

During the cue period, the left and right side light ports flicker based on a Bernoulli process (**Figure 1B**). The odds were set so that, in each 100ms time bin of the cue period, the correct reward port would illuminate briefly (10 ms) with a 75% probability, while the incorrect port had a 25% chance of illuminating. During the cue period the animal was free to respond at any time. In contrast to previous versions of the pulse-based accumulation tasks (Brunton et al. 2013; Scott et al. 2015; Gupta et al. 2023) rats were not required to maintain fixation at the center poke and could move their head freely during the cue period. The trial ended when the rat poked its nose into one of the two side light pokes. If the poke with the higher generative probability was selected, the animal received a drop of sugar water (10% sucrose), if the opposite side was selected the animal received no reward.

We trained Long-Evans rats (n=13; 10 females, 3 males) in this task. Rats were trained in a two-hour session once per day, five days a week. After progressing through a series of training stages (Methods), rats performed an average of 409 trials per session, reached high accuracy (76% correct on average) and exhibited relatively long response times (RT, the time between when the rat initiated its trial and reported its choice) (mean = 0.956s) (**Figure 1C**, **1D**). We observed no differences between males or females in trials per session (One-way RM ANOVA, p=0.458, F=0.592), accuracy (One-way RM ANOVA, p=0.469, F=0.564), or RT (One-way RM ANOVA, p=0.393, F=0.789) .

Consistent with an evidence accumulation strategy, rats exhibited increased accuracy on trials with longer RTs (**Figure 1E**). Next, we sought to evaluate how well the rats’ RT distributions and accuracy were fit by drift diffusion models (**Figure 2**). The traditional DDM provided a good description of the rats’ RT distributions (**Figure 2B**) and indicated relatively high decision bounds, consistent with evidence accumulation. Next, to determine if the model fits could be improved by taking into account the discrete nature of the stimulus we used a pulse-based DDM (pDDM). The pDDM used the same 4 parameters as the DDM, and was similar with the exception that the drift only occurs at stimulus presentation, rather than continuously during the trial (Methods). Both models explained the full distribution of choices and response times equally, determined by comparing the negative log likelihood of the models (Exact Two-Sided KS Test, p=0.9992, **Figure 2C**). Moreover, we did not observe significant differences in the magnitude of side-bias and non-decision time (the time an animal spends not making a decision e.g. movement time etc.), between the DDM and pDDM **(Figure 2D**). The pDDM did consistently predict larger drift rates and boundary parameter values (Exact Two-Sided KS Test, p(drift rate)=1.923e-7, p(boundary)=0.00275, **Figure 2D**), however, these increases were correlated such that both models predicted similar RT distributions (**Figure 2B**). Taken together, these results suggest that rats use an evidence-accumulation strategy to make a decision and that the pDDM and DDM provide an equally good description of rat RTs during the task.

**Figure 2.**
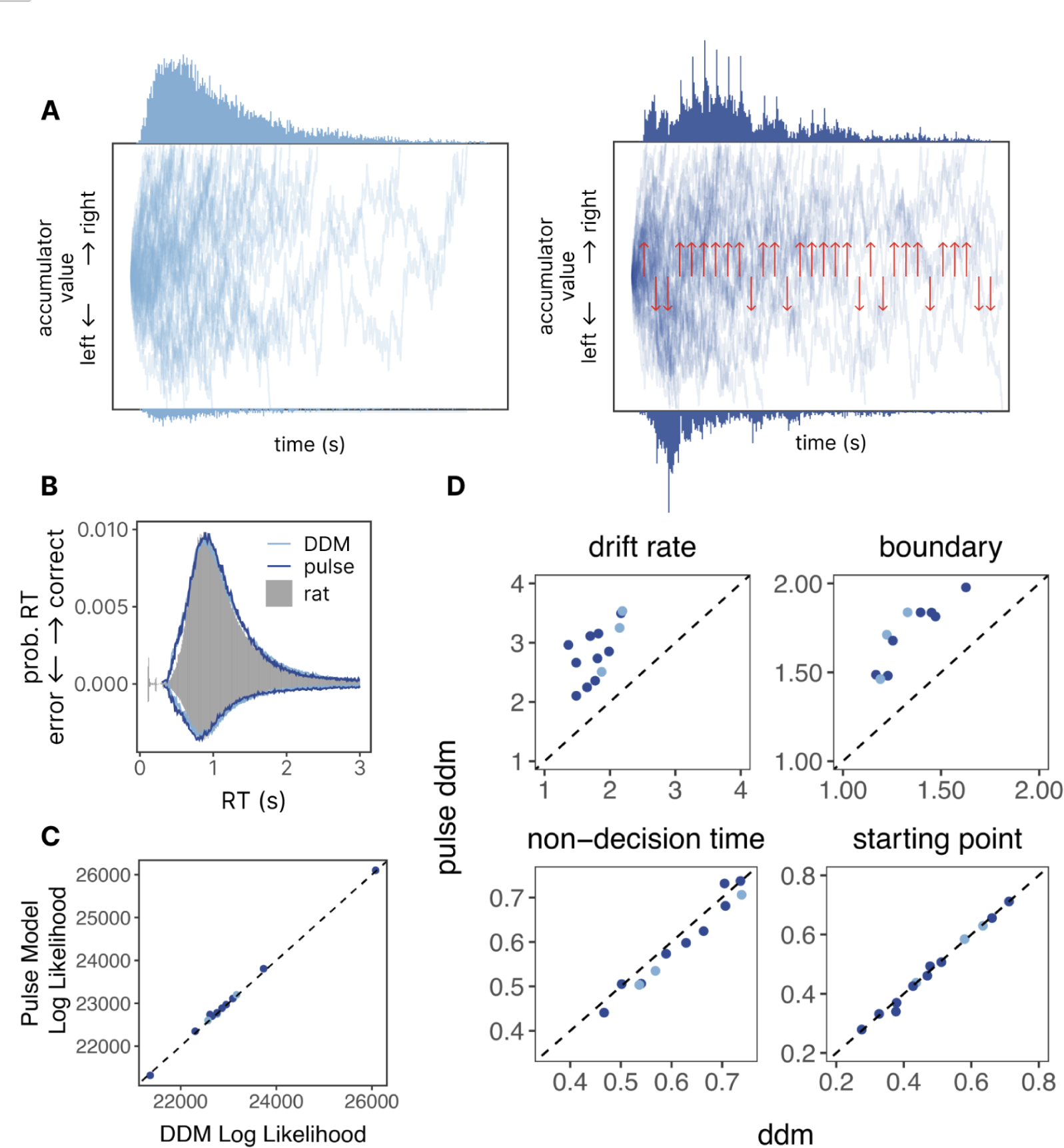
Response times in the accumulation task are well described by diffusion models. A) Schematic of the accumulator value across 50 example trials as predicted by the drift diffusion model (DDM, left panel) and pulse-based model (right panel). The top histogram represents model predicted response times to the upper boundary (i.e., decisions to go right), the bottom histogram represents model predicted response times to the bottom boundary (i.e. decisions to go left). B) Histogram of response times for a single rat showing correspondence between the data (gray), DDM model (light blue line) and pulse model (dark blue line). C) Comparison of the goodness of fit (log likelihood) for the pulse-based model (y-axis) and DDM (x-axis). Each dot represents a different animal. D) Comparison of 4-parameters - drift rate, boundary, non-decision time and starting point - between the pulse-based model (y-axis) and DDM (a-axis). Each dot represents an individual rat. Both models predict similar non-decision times and starting points, whereas the pulse model predicts a larger drift rate and boundary value.

### Rat’s head movements during decision making are well fit by a sinusoidal model

In our task rats are free to move their heads during the cue period. This provided an opportunity to study head movements during evidence accumulation. We recorded video from a subset of rats (N = 6), extracted their movement trajectories, and fit a five parameter sinusoidal model to their head movements on each trial.

A ResNet50-based DeepLabCut network (Mathis et al., 2018) was trained to track the nose, ears, and back of the head of rats as they performed the perceptual-decision-making task (**Figure 3A-B**). The position of the center of the head and the angle of the head relative to the nose poke wall were derived from these key points. To characterize movement trajectories using a small number of interpretable parameters, we utilized a five-parameter sinusoidal model (**Figure 3C**). This model served to represent several aspects of the rat’s behavior, including: i) delay: the time taken to initiate movement, ii) shift: the position of the rat’s head at the start of the trial, and iii) period: the total sinusoidal periods present in the trajectory. The model also incorporated iv) offset: the movement period itself, reflecting head speed through the trajectory, and v) magnitude: the magnitude of the trajectory. The sinusoidal model was fit to the head movements of each rat on each trial and explained most of the data variance with an median R^2 value of 0.9739 **(Figure 3D**). Model fits declined slightly but significantly for longer RT trials (**Figure 3E)**. However the model provided a good description of both correct and error trials for both left and right choices (**Figure 3E**). These results suggest that rats’ head movements during perceptual decision making in a three-port operant chamber are well captured by a 5 parameter sinusoid model.

**Figure 3.**
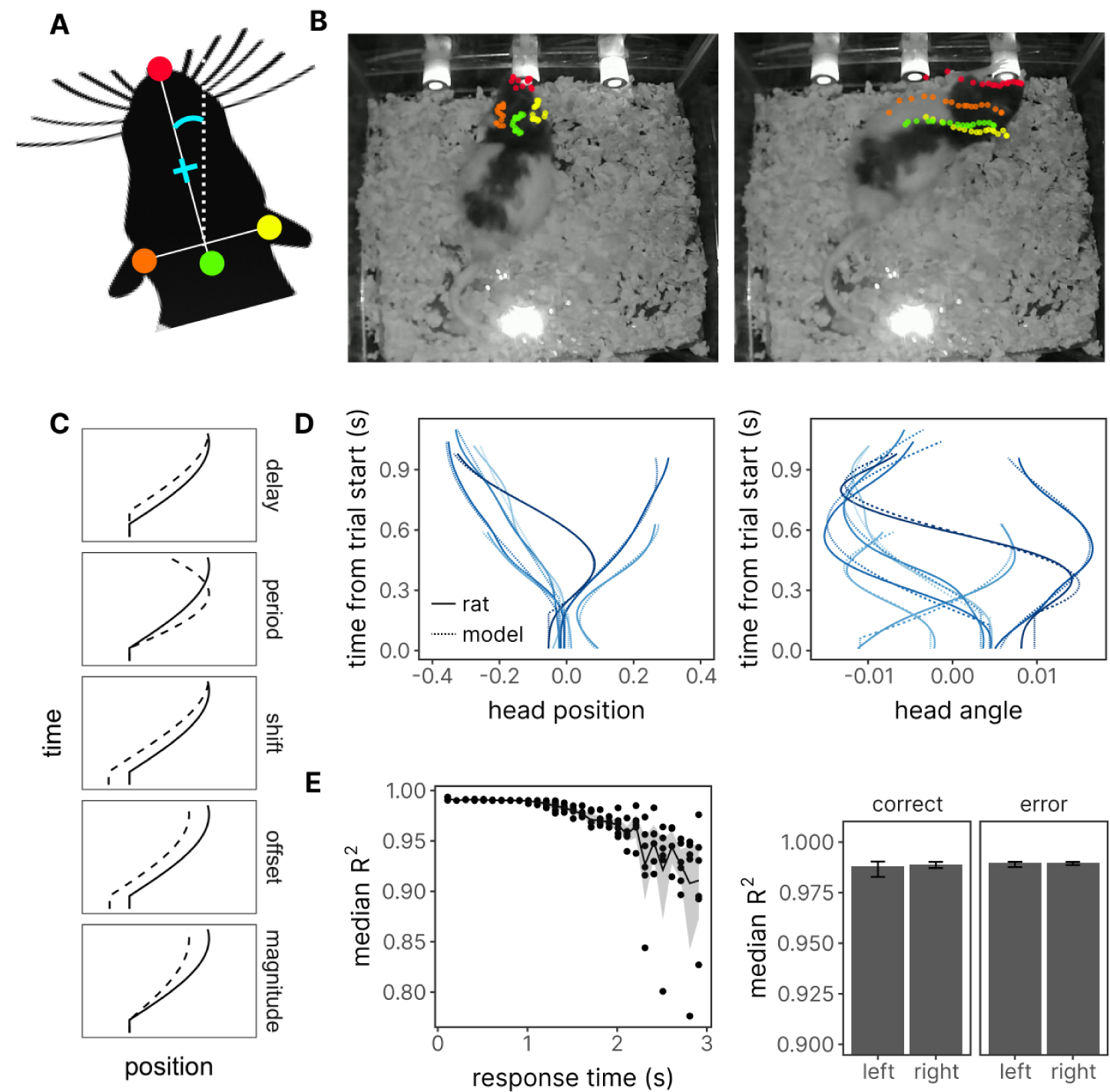
Rats’ head movement trajectories during decision making are well described by a 5 parameter model. A) Schematic of the key tracking points on the rat head. Colored dots represent the four target features tracked by DeepLabCut: nose (red), left ear (orange), right ear (yellow) and back of the head (green). The teal plus sign and angle indicate the center of head and head angle, respectively, derived from these target features. B) Example images from a single trial overlaid with the position of the target features described in panel A. Left panel shows the rat entering the center port with target features for the previous 10 frames (∼300 ms) overlaid. Right panel shows the rat entering the right port with the target features for the previous 10 frames overlaid. Note the rightward trajectory of the animal indicated by the trail of dots reflecting the rightward choice of the animal. C) Schematic of each of the 5-parameters in the head position model. D) Trajectories for head position (left) and angle (right) position on eight example trials. Solid blue lines indicate the data and dashed blue lines indicate model fits. E. Goodness of fit of the model (R^2) across trials with different response times (left) and across correct and error trials.

### Rat’s movements provide information about internal decision variables

A long-standing question in the field is whether movements prior to a decision are reflective of the latent decision making process (Shadlen and Kiani, 2013; Wipinski, 2020). To address this question, we tested whether DDM models with movement information (i.e. mDDMs) could explain rat choices and RTs better than a DDM without movement information (i.e. a traditional DDM). We developed three different mDDMs variants to test three hypotheses about the relationship between movement parameters and decision parameters (**Figure 4**). Our first hypothesis was that, on each trial, the time the rats spent moving to the side-reward port positively correlates with non-decision time. This hypothesis was evaluated using a model where the non-decision time was a function of the movement time (defined as response time minus the delay in the sinusoidal model). We referred to this variant as the Movement Time DDM (mtDDM) (**Figure 4A-B**).

**Figure 4.**
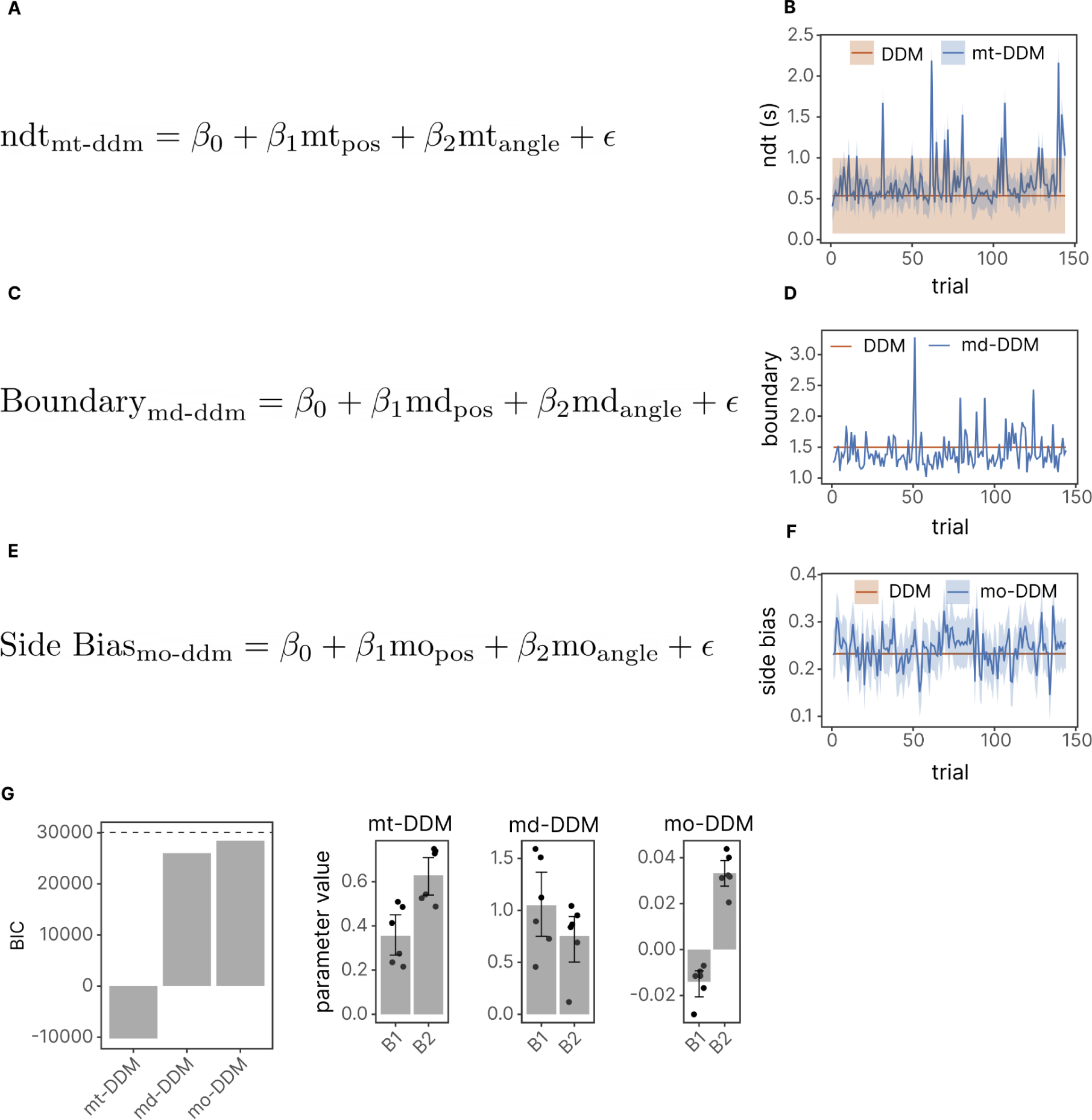
Integration of movement variables improves diffusion model fits. A) Equation for the non-decision time (ndt) in the movement DDM (mt-DDM) model. B) Comparison of the ndt predicted by the DDM and mt-DDM during an example behavioral session. C) Equation for the decision boundary in the movement delay DDM (md-DDM) model. D) Comparison of the boundary predicted by the DDM and md-DDM during an example behavioral session. E) Equation for the side bias in the movement offset DDM (mo-DDM) model. F) Comparison of the side bias predicted by the DDM and mo-DDM during an example behavioral session. G. *Left*: Model comparison (BIC) between the mt-DDM, md-DDM and mo-DDM. Horizontal dashed line represents the performance of the original DDM before incorporating movement parameters. *Right*: Values for each of the two movement parameters (position and angle) in each of the three models, mt-DDM, md-DDM and mo-DDM.

Our second hypothesis postulated that the delay before a movement onset predicts the decision boundary parameter. This hypothesis reflects the intuitive idea that if a rat spends more time waiting before starting to move, it spends more time accumulating evidence. To test this hypothesis, we used a model where the boundary was a linear function of the delay in the sinusoidal movement model. This variant was known as the Movement Delay DDM (mdDDM) (**Figure 4C-D**).

The third hypothesis proposed a correlation between the initial position of a rat’s head and its side bias. For example, if a rat’s head is slightly angled to the right at the start of a trial, it would be more likely to go right than if it were angled slightly to the left. We tested this hypothesis using a DDM variant where the side bias parameter was a function of the side bias as measured by the sinusoidal movement model. We termed this model the Movement Orientation DDM (moDDM) (**Figure 4E-F**).

We compared the performance of each of these new models to the standard DDM. After fitting to rat behavioral data all three movement-informed models (mtDDM, mdDDM, and moDDM) resulted in a lower Bayes Information Criterion (BIC) value compared to the standard DDM **(Figure 4G**). This suggests that the non-decision time variable is the most crucial for improving model fits. This means that rats have significant variability in how much time they take to report their choices after movement onset, which could be indicative of other latent dynamics of the decision process. This suggests that incorporating movement parameters into decision models can improve model fits.

### Rat’s weigh evidence prior to movement onset most heavily

Our model comparison approach suggests that movement time is correlated with non-decision time and that the delay in movement onset is correlated with decision bounds. One intuitive explanation for this result is that the pre-movement period reflects a period of deliberation where the animal weighs sensory information more strongly. To assess this hypothesis, we took advantage of the known timing of sensory pulses and measured the influence of stimuli before and after movement on the animal’s decision using a binomial generalized linear model (GLM) (**Figure 5A)**. We fit three separate models: “All’’, “Before”, “After”. The “All” model represents the differences in right vs left flashes across the entire trial period (“All” model). The “before” and “after” models correspondingly only consider the differences in flashes before movement onset (“Before” model), after movement onset (“After” model), To assess the goodness of fit of each model, we used 10-fold cross validation and used the negative log-likelihood as our objective measurement and compared the predicted psychometric function of each model to the rats’ data. (**Figure 5B-C**). We found that, across all rats, the “All’’ and “Before” model were largely indistinguishable, and were better than the “After” model. These results are consistent with a model in which rats base their decision on an accumulation process in which sensory information received prior to movement onset is weighted most heavily.

**Figure 5.**
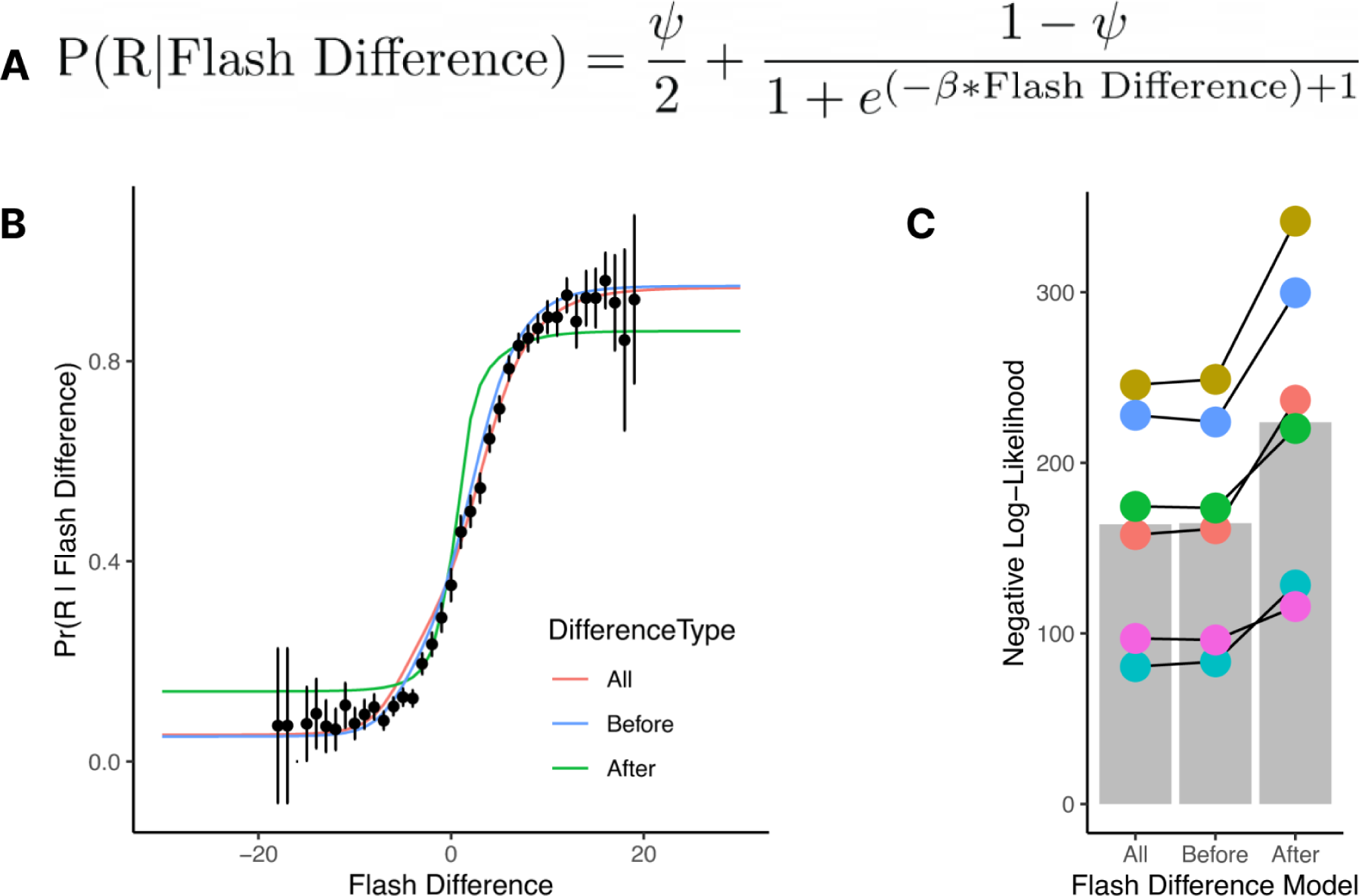
Rats’ choices are better predicted by stimuli before movement onset than stimuli presented during movement execution. A.) Functional form of the model for the movement flashes regression analysis. 𝜓 represents the lapse parameter. B.) Psychometric curves comparing model predictions and rat behavior. Black dots represent rat data, error bars reflect the 95% confidence interval. Colored lines represent the predictions of the model shown in A. In the “All” model flash difference is computed using the entire stimuli presented during the trial. In the Before model, only the stimuli presented prior to movement onset are used. In the “After” model, only stimuli presented during movements are used. C.) Bar plot comparing the negative log-likelihood plots for all fitted models. Colored dots represent model fits for each of the six rats. “Before” and “All” are the best fits to the behavioral data.

## DISCUSSION

In this study, we developed a free response version of a visual pulse-based evidence accumulation task and trained rats to perform this task in a high-throughput operant facility. Behavioral analysis suggests that rats solve this task by perceptual integration: high-accuracy with a positively skewed response time distribution, accuracy increases with response time, choices, and response times were well fit by DDMs. Furthermore, we show that the use of head related positional data to estimate DDM parameters on a trial-by-trial basis (mDDM) provided better fits than a traditional DDM. We also found that even though rats are free to move during the entire trial, rats tend to exhibit a period of fixation prior to movement to their choice.

Interestingly, we find that sensory information presented during this interval prior to movement initiation was as effective in predicting choice as stimuli presented over the whole-trial and significantly better than stimuli presented after movement initiation. Together these findings suggest that this task is useful for studying the decision making process, that rats are capable of performing a free-response task with high accuracy and long response times, and that movement related information can be used to infer the latent parameters of a DDM.

These results are inconsistent with the serial model of decision making, in which motor output follows the decision process (Hanks and Somerfield 2017). For example, our results indicate that head position and angle prior to accumulation predicts the starting point of accumulation in a drift diffusion process. In contrast, our observations are consistent with the parallel or “continuous flow” model, and the embodied cognition model (Lepora and Pezzulo 2015; Kim et al., 2021). Both parallel and embodied cognition models have received substantial support in recent years. In virtual reality navigation-based decision tasks involving perceptual decision, mice typically make pre-emptive movements to a choice target during the accumulation and delay phases of the VR environment (Pinto et al. 2018). Furthermore, rats’ head movement seem to reflect changes of mind (Resulaj et al. 2009) and deliberation, such as vicarious-trial-and-error events at decision points in mazes (Redish 2016). It has also been shown that the eye saccades and reaching behaviors of primates (including humans) reflect internal decision states like indecisiveness and future choices (Seideman, 2018; Król and Król, 2019; Shadmehr et al, 2019; Korbish et al. 2022).

Although our data is consistent with growing evidence for the parallel and embodied cognition models, it is interesting to note that freely moving rats exhibit a phase of relative immobility which corresponds to a period of maximal sensory evidence accumulation. This observation might appear supportive of the serial model or a hybrid model in which animals alternate between brain states where movement and cognition are differently coupled.

### Comparison of diffusion models of decision-making

One of the advantages of the task we report here is that we are able to measure response times which enables fitting of the traditional DDM and a more direct comparison between accumulation models. DDMs are used to delve deeper into the decision-making process, researchers often employ these to model an agent’s behavior in two-alternative forced choice (TAFC) tasks. DDMs assume that an agent continuously accumulates evidence for one option versus another until a level of evidence reaches a predefined threshold, leading to a decision (Ratcliff, 1978, Bogacz et al, 2006). DDMs are instrumental in inferring latent process variables that profoundly affect an observer’s behavioral strategies and have greatly increased our understanding of the evidence accumulation phenomenon (Bogacz et al., 2006).

The data from this task allowed us to fit both traditional DDMs in which the drift occurs continuously throughout the cue period, as well as the pDDM, in which the drift is coupled to the exact time of stimulus presentations. For the data set described here, the continuous and pulse models perform equivalently. However we point out that the pDDM we fit is just one example of a much larger range of accumulation models designed for pulse-based tasks. For example Brunton and colleagues designed a 9-parameter accumulation model which incorporated estimates of noise from a variety of sources (Brunton et al. 2012). Perhaps alternative parameterizations of the pDDM would outperform the DDM in fitting to data from this task.

We chose to use the continuous DDM as a foundation for our movement DDM for several practical reasons. First and most importantly, it provided a good description of choices and response times in this task. Secondly, it was significantly less computationally intensive to calculate the trial-by-trial likelihood than the pulse DDM. Ultimately, this approach revealed that information from rat’s movement trajectories improved inference of DDM parameters (i.e., that the parameters better described rat behavior) on a trial-by-trial basis, as indicated by an improvement in the Bayes Information Criterion (BIC). In future studies it may be possible to incorporate the movement variables into the pDDM which may improve model performance further.

### Comparison with other accumulation tasks

Perceptual decision making is commonly studied using two types of accumulation tasks: continuous and pulse-based. Continuous accumulation tasks present a steady stream of evidence that an observer can integrate continuously. For instance, in the Random Dot Motion (RDM) task, observers must determine the direction of motion encoded by a field of stochastically moving dots which are presented throughout the cue period (Roitman and Shadlen, 2002). In contrast, evidence in pulse based tasks is presented in a stream where the timing of each packet of information is precisely known and readily controllable. This stimulus design facilitates quantitative analysis and precise interrogation of the decision process (Brunton et al., 2013).

Several variants of pulse-based tasks have been used to study evidence accumulation in a variety of species, including humans (Brunton et al., 2013; Keung et al, 2019, Do et al., 2023), rats (Brunton et al. 2013, Scott et al. 2015) and mice (Morcos and Harvey, 2016; Pinto et al., 2018). While these previous pulse-based tasks provided valuable insights into the mechanisms of perceptual decision-making, they often include a fixed accumulation period (Brunton et al. 2013; Scott et al. 2015; Scott et al. 2017, Scott et al; 2018, Colizoli et al, 2018). This poses several challenges, chiefly that DDMs are difficult to apply as they depend on the distribution of response times, which are fixed in a non-free-response version.

Recently, Gupta and colleagues reported an auditory pulse evidence accumulation task that allows for free responses, meaning an observer can respond at any point during the cue period (Gupta et al., 2023). The authors demonstrated that rats could learn this task and used the task to ascertain how history dependence contributes to suboptimal choices in decision making. This newer version effectively addresses the issues of other variants by permitting quantifiable response time distributions (Gupta et al., 2023). However, this task was designed to stop presenting stimuli once an animal breaks its fixation in a nose port, which precludes its ability to answer questions related to accumulation during movement periods. In contrast,the task we describe here poses fewer constraints on the animal movements which allows for greater expression of behaviors.

### Limitations of the current study

Our results suggest that there is a separate stage in the evidence accumulation process, prior to movement, when the rat is heavily integrating sensory evidence. It is worth noting that we cannot effectively rule out the possibility that evidence accumulation occurs over the entire trial.

One reason for this could be that across most of the trials, the flash difference across the pre-trial and the entire trial are highly correlated (Pearson’s r=0.96). This suggests that we cannot fully distinguish between the pre-movement and whole trial flashes in terms of predicting choice. However, since the pre-movement is largely indistinguishable from the model fit of the whole-trial flashes, this would still suggest that stimuli integrated prior to movement onset are most important for determining choice.

A second limitation is that our identification of movement onset is based on the sinusoidal model of head movements. While we show this model to be an accurate description of the rat’s head trajectory during our task, we did not evaluate other forms of movement, such as limb or trunk position. It is possible that incorporation of other forms of movement into the DDM would further improve model fits. Body movements and changes in posture can drive changes in circuits that are of interest in evidence accumulation studies, such as the posterior parietal cortex (Mimica et al. 2018; Tombaz et al. 2020). Therefore a more global picture of movements during evidence accumulation may be valuable to better interpret neural dynamics associated with decision making.

A third limitation is that we were not able to determine whether the strength of evidence differs between the pre-movement and movement period. In future studies our approach could be used to quantify the weight of evidence pulses during movement and non-movement phases. Or alternatively, using real time pose tracking methods, closed-loop experiments could be designed to alter evidence during the different phases of the task to probe the influence on the choice process.

### Integration with neural recordings

Prior studies have performed electrophysiological or calcium recording while rodents and primates perform continuous and pulsed-based accumulation tasks (Roitman and Shadlen, 2002; Hanks et al. 2015; Scott et al. 2017; Pinto et al., 2018; Boyd-Meredith et al., 2022). These studies have identified many cells that exhibit changes in firing rate or that correlated with decision and task variables, including the instantaneous evidence and accumulated evidence (or decision variable). Some have interpreted these dynamics as evidence in favor of a neural implementation of the drift-diffusion like process (Gold and Shadlen 2007, Brody and Hanks 2016). However, despite technological advancements in neural measurement and analysis, concrete interpretations of dynamics recorded during these decision tasks have remained elusive (Costello et al., 2013; Musall et al. 2019). However, it is evident that movement is widely represented across the cortex and therefore should be considered as a factor driving neural dynamics during decision making (Musall et al. 2019; Umeda et al. 2019). This presents a difficulty in interpreting neural dynamics recorded during these tasks: to what extent are signals that represent accumulation and decision, separate from movement? Our results suggest that there appears to be a stage of evidence accumulation prior to movement onset, at least in rats performing the visual pulse-based task. Thus, by incorporating video recordings of behavior into decision making models, future studies can better separate neural dynamics related to evidence accumulation from those associated with movement.

### Extension to other species

In this study we show that rats are able to learn the response-time visual pulse-based task in a computer-controlled facility, reach peak performance in a relatively short time and exhibit several hallmarks of evidence accumulation with relatively low lapse rates (16.3%). In future studies it would be useful to extend this task to other species in order to evaluate within and across species differences in evidence accumulation. Previous work has shown that fixed duration visual pulse-based tasks, similar to those developed for rats (Scott et al. 2015), can be performed by mice in a virtual navigation setting (Morcos and Harvey 2016, Pinto et al. 2018) and by humans in a lab setting (Brunton et al. 2013) or as an online game (Do et al. 2023).

Furthermore, non-verbal feedback-based training pipelines, similar to those used in animal studies, have also been incorporated into human studies (Do et al. 2023). These non-verbal training pipelines allow a more direct comparison between human and animal subjects and may allow measurement of evidence accumulation in individuals with difficulty following language-based instructions, such as minimally verbal autistic individuals.

## MATERIALS AND METHODS

All experiments and procedures were performed in accordance with protocols approved by the Boston University Animal Care and Use Committee. Long evans adult rats (N=31; aged 3 months to 2 years) were purchased from Taconic or bred in house. Both male and female rats were used and trained in the same room at the same time. Rats were food restricted to 90-100% of their body weight and fed once per day (typically 3-4 pellets of food per day). They received 0.025-0.04 ml (depending on the session and rat) of 10% sucrose (100g/L). Reward volume was consistent within a session. Both 24-hour trained rats and daily trained rats were housed on a 14:10 ON:OFF light schedule with the ON phase corresponding to daylight hours in Boston, MA USA.

### Behavioral control system

Behavioral control system was inspired by Poddar et al. 2013 and Dhawale et al. 2015. Rats were trained in custom acrylic chambers with three nose ports. Nose ports were 3D printed (Sanworks or custom made) and equipped with a visible LED for stimulus delivery (Sanworks), peristaltic pump for reward delivery, and an IR LED and photodetector as a beam break (Sanworks). Behavioral control software to implement the task and control individual boxes was written in MATLAB. Boxes were controlled through a Teensy-based microcontroller system (Bpod Sanworks). A custom written python application was used to control multiple Bpod instances from a single control computer, requiring an edited version of the Bpod MATLAB software library (edited Bpod library: https://github.com/RatAcad/Bpod_Gen2; Custom python application: https://github.com/RatAcad/BpodAcademy). Further details about the software implementation can be found in the respective code repositories. The floor of the chambers contained bedding with bedding (part no) which was changed after each session.

### Daily training

Rats (n=13) were housed in pairs in an animal facility and moved to the training room in the laboratory each morning for training. Rats ran for 2 hour shifts, 5 days per week in the late morning (10am-noon) or early afternoon (1pm-3pm). Feeding was conducted in the late afternoon after behavioral training (1-4pm). Each chamber also contained a video camera mounted above the chamber.

### Training pipeline

Rats were rewarded with 10% sucrose at all stages of training and testing. Training took 1-4 weeks depending on the rat and progressed through 3 stages. In the first stage (1-3 days) rats were rewarded for inserting their nose into a side port with an illuminated LED. In stage 3 (1-21 days) rats received reward for inserting their nose into the center port and then the side port with an illuminated LED. In the third stage, they received a reward for inserting their nose into the center port and then the side port with a flashing LED. Once rats reached criterion 80-100% correct, the probability of flashes on the incorrect side increased in the following way 100:0 -> 90:10 -> 80:20. Progression through this third stage took 1-2 days per condition. After completion of all stages and conditions rats performed the task with a flash probability of 75:25.

### Behavioral task

At the start of a trial, the light in the center port turned on indicating that the rat could initiate the trial at its leisure. Once the rat nose poked in the center port, a single flash would occur simultaneously in both the left and right ports. After this one light flash on both sides, the rat would see a series of light flashes according to a Bernoulli process – every 100 ms, the rat would see a flash on either the right or the left side with a probability of 75:25 that the flash would occur on the correct side vs. the incorrect side; the correct side was drawn randomly on each trial. To record a response, the rat nose poked in either the left or right port and the series of light flashes would terminate as soon as this decision was recorded. If the rat responded correctly, the light in the correct side port turned on for 2 s and a 30uL sucrose water reward was delivered immediately. If the rat responded incorrectly, all lights turned off and no reward was delivered, and new trials would start after a 2s delay. For daily testing, rats participated in 2 hour sessions Monday-Friday.

To determine if rats displayed a speed accuracy trade off, a generalized additive mixed model was fit to the binary trial outcome (i.e. 1 for correct decision) against RT. We fit trials up to 3s of response time because greater than 90% of trials occurred within this period.

To determine how trial lengths affected the model fit, the trial R^2^ for the observed trajectory was modeled using a mixed effects beta regression with a logit link function to evaluate the model fit as a function of response time and an intercept term. The trial length was found to be statistically significant in predicting the model R^2 (p<2e-16).

### Diffusion models of decision-making

Drift diffusion models (DDMs) are commonly used to describe decision-making in two-choice tasks. DDMs assume that a decision-maker accumulates noisy evidence for one response vs. the other time and commits to a response when the level of evidence reaches a pre-defined threshold. Two diffusion models were applied to behavioral data: the extended drift diffusion model (Ratcliff 1978) and novel pulsed evidence drift diffusion model based on Brunton et al., 2012.

The extended drift diffusion model assumes that, on average, evidence for one option vs. the other grows linearly over time: the change in evidence, *dx*, is updated by a constant drift rate, *v*, and a gaussian noise term, *cdW*:

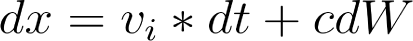

In the extended drift diffusion A decision is made at the first time point at which the evidence *x*, crosses a boundary, *a*. The response time on the decision is then the time from the trial start to the boundary crossing + a non-decision time parameter, *ndt*, which represents any additional time that goes into executing a response, such as the time it takes to move to the reward port after committing to a decision. The extended drift diffusion also includes one additional parameter: the side bias, *x_0_*, which accounts for a bias to go left vs. right – according to the drift diffusion model, a bias to go left means that the animal requires less evidence to commit to the choice to go left vs. right.

The extended drift diffusion model also allows for random trial-to-trial variability in the drift rate, the non-decision time, and the starting point:

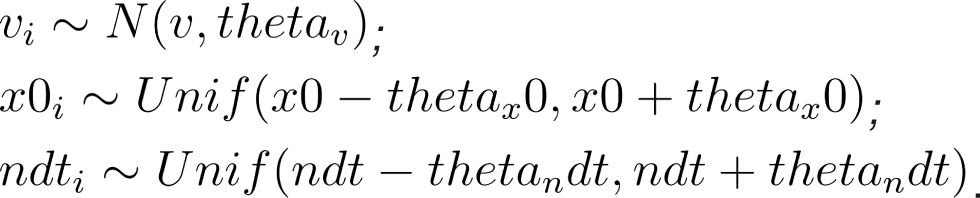

In the pulse diffusion model, evidence is accumulated per unit of stimulus rather than continuously over time: *dx* = *v_t_* * *s_t_* * *dt* + *cdW*

Where *s_t_* is the stimulus at time *t*, *s_t_* during a right pulse, *s_t_* = −1 during a left pulse, and *s_t_* = 0 otherwise.

Additionally, consistent with the pulsed evidence accumulation model by Brunton et al., 2012, the drift rate does not vary trial-by-trial but it drifts over the course of the trial: *v_t_* ∼ *N* (*v, theta_v_*). Furthermore, in this model, variability in the side bias across trials follows a normal distribution rather than a uniform distribution: *x*_0*i*_ *N* (*x*0, *theta_x_*0).

### Video recording and image processing

Video was recorded during a subset of daily testing sessions using USB webcams. Videos were recorded at 30 fps using the same custom python application used to control behavioral boxes(<u>github.com/RatAcad/BpodAcademy</u>). To synchronize video data with behavioral data, the time of each camera frame and a TTL signal indicating the start of the trial (center light turned on) and trial initiation (nose poke in center port) were both recorded using the python *time* package. TTL signals were sent from the Bpod State Machine and recorded using serial input from a Teensy 3.2 microcontroller.

The position of the rat during a behavioral session was extracted from the video data using DeepLabCut (Mathis et al, 2018). Experimenters identified the position of the ears, nose and back of the head (see Figure 3A) on 620 frames across 5 animals which were used as training data (95% of frames) and validation (5% of frames). A ResNet-50 base neural network was used for 160,000 training iterations with a test error of 6 pixels (640-480 pixel images). A p-cutoff of 0.75 was used to condition (X, Y) coordinates for future analysis.

To standardize rats’ positions across sessions despite slight differences in camera position and angle in different boxes, key points were first corrected for any translation and rotation using the following procedure:

1. The (X, Y) coordinates of each nosepokes were defined as the average coordinate of the nose at the time of each nosepoke (e.g., to find the center nosepoke, the average of the nose position on frames that aligned in time with rats’ nose poke in the center port).
2. All keypoints were translated such that the center nose poke was at coordinate (0,0)
3. Keypoints were rotated and scaled such that the left and right nose poke positions were at coordinates (−1, 0) and (1, 0) respectively.

Finally, two keypoints of interest were computed: i) the center of the head as the center point in between the nose, back of head, and between the ears and ii) the angle of the head relative to the nose-poke wall as the angle of the intersection between a line drawn from the nose and back of head and a line drawn between the nose pokes).

### Sinusoidal Movement Model

To reduce the dimensionality of movement trajectories (∼100-300 observations per trial), a sinusoidal model was fit separately to the position of the head and the angle of the head for each rat on each trial. The model was formulated as:

time_of_movement = {

time_from_initation > *t_i* : time_from_initiation - *t_i*

time_from_initiation <= *t_i* : 0

}

predicted_trajectory = choice * *y_s* * sin(*x_s* * time_from_movement + *x_t* / dt) + *y_t*

where time_from_initiation is the time since the rat nose poked in the center port, choice is the rat’s choice on the trial (coded as -1 for left, 1 for right), dt = 0.01 is the resolution of the trajectories, and the following 5 free parameters:

1. *t_i* (movement delay): the time from the center nose poke to the initiation of movement
2. *y_t* (position offset): the lateral position (or angle) of the rat’s head relative to the center nosepoke (i.e., is the rat leaning to the left or right at the start of the trial)
3. *y_s* (movement magnitude): the amplitude of the sinusoidal curve
4. *x_t* (starting phase): the phase of the sinusoidal curve when the rat initiates movement (i.e., is the rat at the beginning of the phase – moving in the opposite direction before turning back – or the end of the phase – moving straight to the chosen reward port)
5. *x_s* (the sinusoidal period): how fast the rat moves through the phase (i.e., did the rat accelerate quickly throughout the movement or progress more slowly)

Free parameters were estimated by maximizing the R^2^ between the predicted and observed trajectories using generalized simulated annealing (GenSA R package; Xiang et al., 2013).

### Data analysis and model fitting

Data analysis was performed in R version 4.3.1. Calculation of standard statistical tests(e.g. KS tests, ANOVAS) were performed using the stats and aov packages. To fit the accuracy vs. response time curve in Figure 1E, we used the mgcv (Wood, 2017) package to fit a generalized additive mixed model. The beta regression was fitted using the glmmTMB (Brooks et al, 2017) package.

DDM and pulse DDM models were fit using a custom R package (rddm; https://github.com/gkane26/rddm). For comparison of the DDM and the pulse DDM, parameters were estimated by maximizing the quantile maximum probability estimate (Heathcote et al., 2002; 2004) with differential evolution as the optimization routine (RcppDE package). To estimate the influence of movement parameters from the sinusoidal model on trial-by-trial changes in DDM parameters, DDM model parameters were estimated using maximum likelihood estimation. Trial-by-trial likelihoods (as implemented in the rddm package) were calculated using the rtdists R package (Singmann et al. 2022; Voss & Voss, 2008).

To fit our lapse parameter binomial GLM we wrote a custom link function in R and performed constrained optimization of the negative log-likelihood function using the L-BFGS-B algorithm in the optim function in R such that the lapse parameter was bounded between 0 and 1. For the optimization we used 10-fold-cross validation. To do this, we fit the model to each rat, and did CV within each animal’s trial data.

All plots were generated using ggplot2.

## ACKNOWLEDGEMENTS

We thank Bence Ölveczky (Harvard University) for sharing details of training chambers and Josh Saunders (Sanworks), Dustin Clark (BU Research Computing Services) and Quan Do for technical assistance with implementing data collection. We thank Sinead O’Brien-Wernig and Amber Hickey and the BU Animal Science Center staff for their care of the animals. We thank Honjie Xia, Arula Ratnakar, Shiping Li, Rifqi Affan and other members of the lab for help with data collection and maintenance of the behavioral training facility. Brian Depasquale and Chandramouli Chandrasekaran provided valuable feedback on the manuscript. BBS was supported by a NIH R56MH132732 Award and a Whitehall Foundation Research Grant. GAK was supported by a Boston University Center for Systems Neuroscience Distinguished Fellow award.

## Author Contributions

Conceptualization: G.A.K. and B.B.S.; data collection: G.A.K; modeling and analysis: G.A.K. and R.A.S; writing – original draft: G.A.K and R.A.S; writing – review & editing: G.A.K and R.A.S. and B.B.S.

